# A depth map of visual space in the primary visual cortex

**DOI:** 10.1101/2024.09.27.615442

**Authors:** Yiran He, Antonio Colas Nieto, Antonin Blot, Petr Znamenskiy

## Abstract

Depth perception is essential for visually-guided behavior. Computer vision algorithms use depth maps to encode distances in three-dimensional scenes but it is unknown whether such depth maps are generated by animal visual systems. To answer this question, we focused on motion parallax, a depth cue relying on visual motion resulting from movement of the observer. As neurons in the mouse primary visual cortex (V1) are broadly modulated by locomotion, we hypothesized that they may integrate vision- and locomotion-related signals to estimate depth from motion parallax. Using recordings in a three-dimensional virtual reality environment, we found that conjunctive coding of visual and self-motion speeds gave rise to depth-selective neuronal responses. Depth-selective neurons could be characterized by three-dimensional receptive fields, responding to specific virtual depths and retinotopic locations. Neurons tuned to a broad range of virtual depths were found across all sampled retinotopic locations, showing that motion parallax generates a depth map of visual space in V1.

## Introduction

To guide behavior using vision, the visual system must infer the three-dimensional location of visual cues based on the two-dimensional images formed on the retinae. The importance of this computation is underscored by the fact that the capacity for depth perception is innate in many animals (1). However, our understanding of how the visual system parses the organization of three-dimensional visual scenes to infer the depth of visual cues remains limited. Computer vision algorithms for depth estimation use two-dimensional input images to generate depth maps of the visual scene, where every pixel represents the estimate of the depth at that location (2). Such explicit representations of depth help guide the behavior of artificial agents (3). Do animal visual systems also generate depth maps of the visual scene and how are such depth maps encoded in neural activity?

In mammals, depth perception is supported by multiple cues relying on both binocular and monocular information (4–9). While binocular vision supports fine depth judgments (10), experiments in animals and humans show that it is not necessary for depth perception (1, 11–13). In humans, motion parallax, a monocular depth cue that relies on visual motion arising from movement of the observer in the environment, is sufficient to generate a sensation of depth in the absence of other depth cues (14). When animals move, nearby and far visual cues appear to move at different velocities (15) making it possible to estimate their depth based on the speed of movement of the observer and the rate of the resulting visual motion. Previous work in passively translated macaques has found evidence for representation of depth from motion parallax in area MT (6, 16, 17). However, little is known about how visual circuits in mammals process motion parallax resulting from active movements of the observer and how the resulting representations are mapped across the visual field.

In mammals with limited binocular overlap, such as rodents, motion parallax is thought to be one of the major sources of depth information (12, 13). In mice, eye movements stabilize the direction of gaze during locomotion to compensate for head rotation (18–20). Consequently, self-generated visual motion is primarily driven by translation of the animal, and its speed is proportional to the ratio of locomotion speed and depth (Figure 1A). When performing tasks that require depth perception, mice can make monocular depth judgments that depend on the intact activity of the primary visual cortex (V1) (13), but how V1 neurons support depth perception from monocular cues such as motion parallax is unknown. Visual responses in mouse V1 are broadly modulated by locomotion (21–23) and the functional role of this modulation in visual processing remains debated (24, 25). As estimation of depth from motion parallax requires both visual and self-motion information, we hypothesized that it is supported by locomotion-related modulation of visual responses in V1.

**Figure 1.**
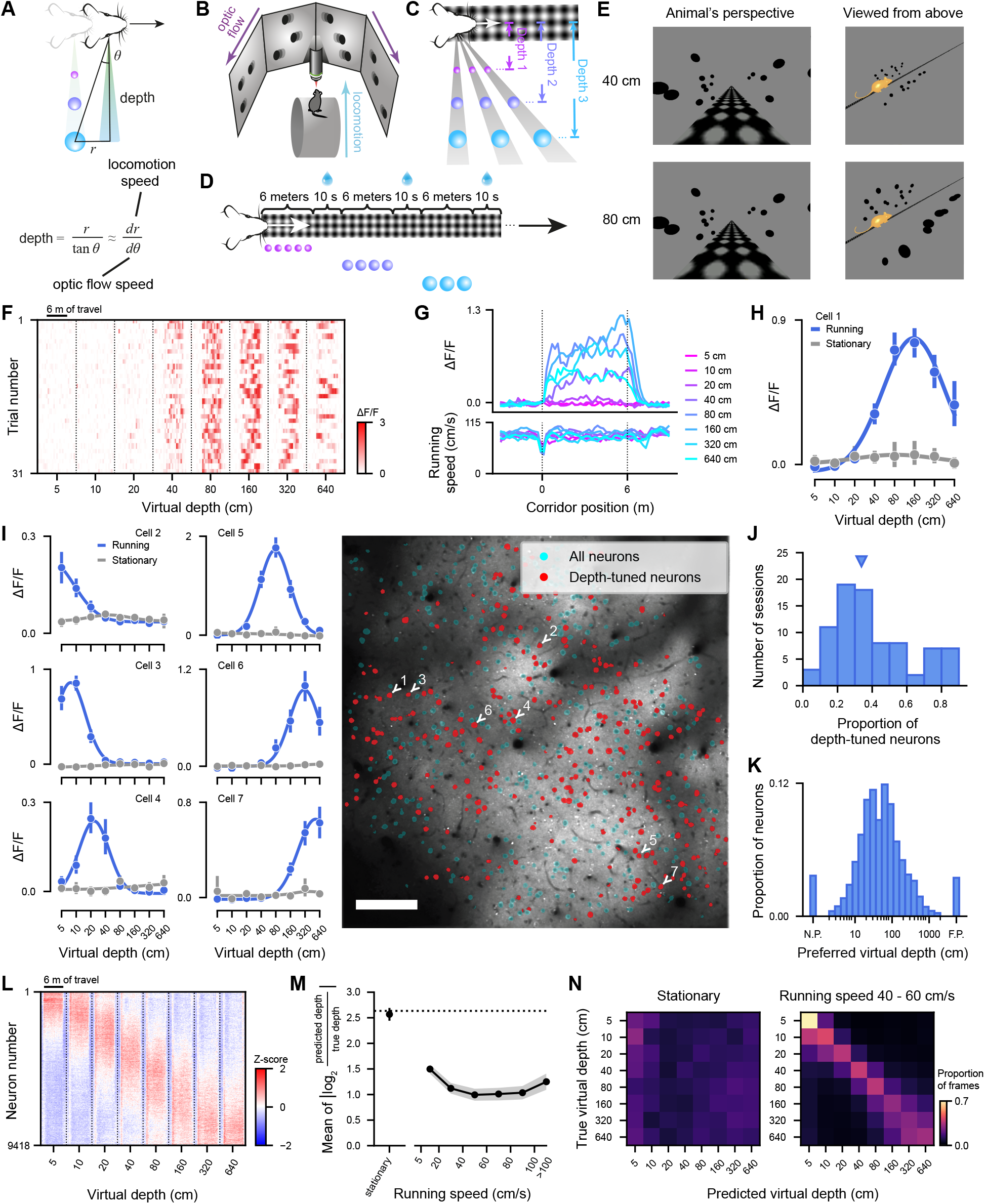
Depth selectivity from motion parallax in layer 2/3 of mouse primary visual cortex. **(A)** Schematic of the relationship between locomotion and visual motion speed due to motion parallax. **(B-C)** Virtual reality setup and stimulus schematic. **(D)** Schematic of trial structure. **(E)** Illustration of stimuli at two virtual depths from the animal’s perspective and viewed from above. The mouse (not to scale) is shown to indicate the animals position in the virtual environment. **(F)** Raster plot of responses of an example neuron in mouse V1 across virtual depths as a function of distance traveled during the trial including 3 m of inter-stimulus interval before and after each trial. Dashed lines separate different trial types. **(G)** Mean responses of the example neuron (top) and mean running speed (bottom) across virtual depths. **(H)** Virtual depth tuning of the neuron in **F-G** during locomotion (>5 cm/s) or stationary (max speed during preceding 1 second <5 cm/s) periods. Error bar – 95% confidence interval. **(I)** Virtual depth tuning of 6 additional example neurons during running (blue) and stationary (gray) periods (left) and spatial location of depth-tuned neurons in an example imaging session (right). Scale bar – 100 *µ*m. Error bar – 95% confidence interval. **(J)** Proportion of depth-tuned neurons across imaging sessions (N = 85 sessions). Triangle indicates the median value. **(K)** Distribution of preferred virtual depths of depth-selective neurons (N = 31,013 neurons). N.P., near-preferring neurons; F.P., far-preferring neurons. **(L)** Raster plot of z-scored responses of all depth-selective neurons from sessions with 8 virtual depths in mouse V1, in a [-1 m, +1 m] window around stimulus presentation. Neurons are sorted by virtual depth preferences estimated using hold-out data not used in calculation of the raster plot. **(M)** Mean error of virtual depth decoded from population activity of simultaneously recorded neurons when mice were stationary and as a function of running speed (N = 85 sessions). Dashed line – chance level; shading – 95% confidence interval. **(N)** Average confusion matrices of decoded virtual depth when animals were stationary or running at a speed of 40-60 cm/s (N = 60 sessions with 8 virtual depths).

To test this hypothesis, we designed a virtual reality environment for head-fixed mice, where motion parallax acted as a cue of virtual depth, and recorded the activity of excitatory neurons in layer 2/3 of V1 using two-photon calcium imaging as mice navigated through this environment. We found that selectivity for virtual depth from motion parallax was widespread among V1 neurons. This depth selectivity could not be explained by visual-evoked responses alone but resulted from the integration of optic flow and locomotion-related signals.

Furthermore, by reconstructing visual stimuli presented in the virtual environment, we mapped the three-dimensional receptive fields of V1 neurons – tuned for both retinotopic location and virtual depth of visual cues. By characterizing depth selectivity across V1, we show that V1 neurons responding to stimuli across all sampled visual field locations encode the full range of virtual depths indicating that motion parallax generates a depth map of visual space in the visual cortex. However, depth selectivity was not homogeneously distributed across V1, with regions representing the upper visual field enriched for neurons tuned to near depths. We speculate that this bias reflects the ethological significance of the upper visual field for detection of nearby threats.

## Results

### Depth selectivity from motion parallax in virtual reality

To test whether neurons in mouse primary visual cortex (V1) integrate visual- and locomotion-related signals to estimate depth from motion parallax, we trained head-fixed mice to navigate a virtual reality (VR) environment where visual stimuli were presented at different virtual distances from the mouse and motion parallax acted as a cue of depth (Figure 1B-E). The environment contained a floor texture that the mouse ran across and black spheres presented in the monocular visual field (Figure 1B,E). The position of the mouse in the VR environment was updated in closed loop based on its running speed on a styrofoam cylinder (Figure S1). Virtual depth of spheres varied from trial to trial (logarithmically spaced, with either 5 depths ranging from 6 cm to 600 cm in 25 sessions, or 8 depths ranging from 5 cm to 640 cm in 60 sessions). Consequently, animals’ locomotion on the wheel generated different speeds of optic flow depending on the virtual depth on a given trial. New spheres were presented in front of the mice as they advanced through the virtual environment to provide constant visual stimulation throughout the trial. Each trial was terminated once mice traveled 6 m through the VR, after which the spheres disappeared. To motivate mice to run, they were food-restricted and received a probabilistic soy milk reward after the spheres disappeared on 60-80% of trials (Figure 1D). The size and density of spheres on each trial were adjusted such that spheres subtended the same visual angle across depths and had similar spacing from the animals’ perspective (Figure 1C). The animals running speed was similar across trials of different depths (Figure S2A-D) and the sphere stimulus did not elicit any stereotyped eye movements (Figure S2I-L).

We recorded the activity of neurons in layer 2/3 of V1 in transgenic mice expressing GCaMP6f or GCaMP6s in excitatory cortical neurons using two-photon calcium imaging (60,357 neurons from 85 sessions in 7 mice) and determined whether neuronal responses were selective for virtual depth. To this end, we quantified virtual depth tuning by calculating the mean calcium fluorescence for trials of different virtual depths (Figure 1F-H). Depth selectivity was widespread in among layer 2/3 excitatory cells, with individual neurons tuned to different depths (Figure 1I-J). Across the population, 51.4% of cells (31,013 of 60,357 neurons) exhibited significant depth selectivity and virtual depth preferences of depth-selective neurons spanned the full range of depth values that we tested (Figure 1K-L).

Depth-selective responses were absent during periods when mice were stationary (maximum speed during preceding 1 second <5 cm/s, Figure 1H-I). At the population level, virtual depth could be decoded from V1 population activity during locomotion but not during stationary periods (Figure 1M-N). This demonstrates that motion parallax resulting from animals’ locomotion gives rise to a representation of depth in mouse V1.

To ensure that sphere stimuli subtended the same visual angle across different virtual depths, their size in the virtual environment was proportional to depth on a given trial (Figure 1C). Therefore, depth selectivity could in fact reflect selectivity for virtual size. To test this possibility, we simultaneously varied virtual depth and virtual size of the spheres in a subset of recordings, such that the spheres subtended either 5, 10 or 20 degrees of visual angle (Figure S3). The magnitude of neuronal responses was modulated by the visual angle of the spheres (Figure S3B), consistent with the widespread size tuning of V1 responses (26, 27). However, the preferred virtual depth was consistent across trials with 5, 10, and 20 degree spheres (Figure S3B-C), demonstrating that selectivity for virtual depth is invariant to stimulus size.

### Depth selectivity arises from the integration of optic flow and locomotion-related signals

Statistics of optic flow speed differed across virtual depths – near depths generate faster optic flow than far depths (Figure S2E-H). Consequently, depth selective responses could arise solely as the product of selectivity for optic flow speed. On the other hand, as the speed of self-generated optic flow depends on the ratio of the speed of locomotion and virtual depth, depth-selective neurons may preferentially respond to the conjunction of these variables.

To determine how the depth selectivity arose as a function of optic flow and locomotion speeds, we calculated the optic flow speed for each imaging frame as the ratio of running speed and virtual depth, which corresponds to the rate of visual motion at 90 degrees azimuth. We then visualized the activity of depth-selective neurons as a function of running speed and optic flow speed (Figure 2A-B, S4A-B,E-F). If optic flow responses were the only determinant of depth selectivity, optic flow speed tuning should not change across virtual depths. However, this analysis showed that depth selectivity persisted when controlling for the speed of optic flow (Figure 2B, S4B,F). As mice have to run at different speeds to generate the same rate of optic flow across different depths, this supported the hypothesis that depth-selectivity arose from the interaction of optic flow and locomotion-related signals. To examine this directly, we analyzed neuronal responses as a function of both optic flow and running speeds. This analysis revealed that depth-selective neurons in V1 responded to a specific conjunction of running speed and optic flow speed (Figure 2C, S4C,G).

**Figure 2.**
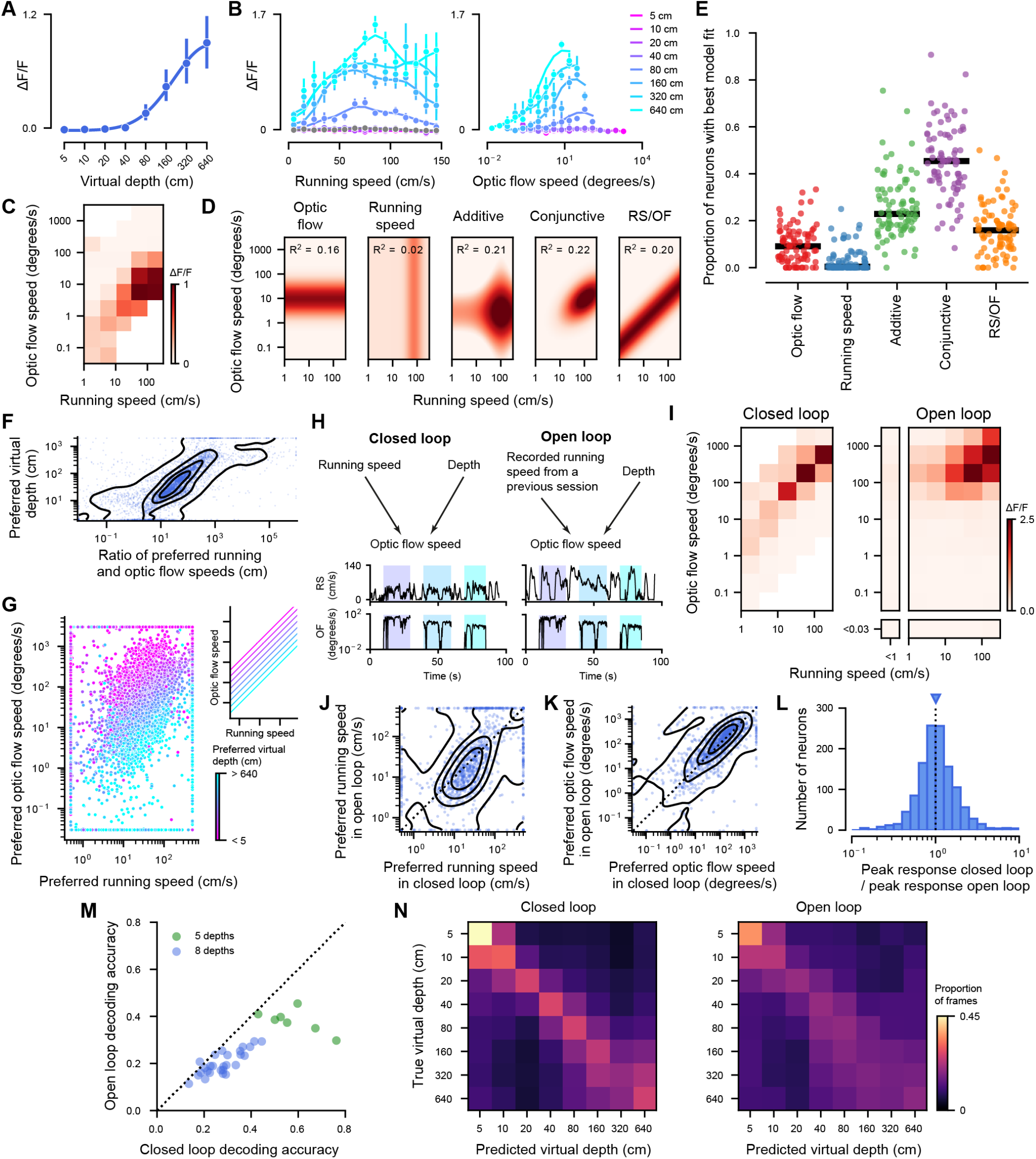
Depth selectivity arises from conjunctive coding of optic flow speed and running speed. **(A-B)** Virtual depth tuning **(A)** and responses of an example depth-selective neuron binned by running speed **(B** left) or optic flow speed **(B** right). Gray line **(B** left) – running speed tuning during the inter-trial interval, when sphere stimuli were absent. **(C)** Responses of the example neuron in **A-B** and as a function of both running and optic flow speeds. **(D)** Running speed and optic flow tuning of the example neuron in **A-C** fitted with the pure optic flow, pure running speed, additive, conjunctive or ratio models (see methods). RS – running speed, OF – optic flow speed. (**E**)Proportion of depth-selective neurons best explained by the pure optic flow, pure running speed, additive, conjunctive and ratio models in each session (N = 85 sessions). (**F**)Preferred virtual depths as a function of the ratio between preferred running and optic flow speeds estimated using hold-out data not used in calculation of preferred virtual depths (N = 5,737 neurons). Contours – kernel density estimate. **(G)** Preferred optic flow speed and running speed of neurons estimated using hold-out data color coded by their preferred virtual depth (N = 5,737 neurons). Inset – optic flow speed as a function of running speed across virtual depths. **(H)** Relationship between running speed and optic flow speed on closed loop and open loop trials and example traces. Shaded areas color-coded by depth indicate stimulus presentation. **(I)** Responses of an example neuron as a function of both running and optic flow speeds in closed loop (left) and open loop (right). **(J-K)** Scatter plot of preferred running speed **(J)**, preferred optic flow speed **(K)** in open loop and closed loop for depth-selective neurons (N = 1,234 neurons). Contours – kernel density estimates. **(L)** Histogram of ratios of peak response amplitude in closed loop over open loop for neurons in **J-K. (M)** Accuracy of decoding virtual depth from population activity of depth-selective neurons in closed loop and open loop recordings (N = 34 sessions). **(N)** Confusion matrices of decoded virtual depth in closed loop and open loop recordings with 8 virtual depths (N = 27 sessions).

To quantify how running speed and optic flow speed tuning were integrated, we fitted neuronal responses as a function of running speed and optic flow speed using five classes of models: models tuned to either optic flow or running speed in isolation; an additive model, which models neuronal responses as linear summation of optic flow and locomotion-related responses; a conjunctive model, where neurons are tuned to a specific conjunction of optic flow and running speeds defined by a two-dimensional Gaussian function; and an idealized depth tuning model, where neuronal responses depend on the ratio of locomotion and optic flow speed (Figure 2D, S4D,H). To control for differences in model complexity, we used K-fold cross-validation to assess the performance of each of the 5 models on test data not used for parameter optimization. We found that the pure optic flow and running speed models could not explain neuronal responses, showing that integration of visuo-motor signals is required to account for depth selectivity. The conjunctive model consistently outperformed the other models (Figure 2E; *p* < 0.0001 vs optic flow model; *p* < 0.0001 vs running model; *p* < 0.0001 vs additive model; *p* < 0.0001 vs ratio model). Therefore, responses of depth selective neurons are best explained by selectivity for a specific combination of running and optic flow speeds.

We next asked how the preferred virtual depth of individual neurons related to their preferred optic flow and running speeds. To this end, we split the trials recorded for each neuron into two sets: one half of the trials was used to estimate each neuron’s preferred virtual depth, while the other half was used to estimate preferred optic flow and running speeds using the conjunctive model described above. Due to motion parallax, the ratio of running speed and the resulting optic flow is equal to virtual depth (Figure 1A). Consistent with this, we found that the ratio of the preferred running and optic flow speeds of individual V1 neurons could predict their preferred virtual depth estimated using hold out data (*r* = 0.794, *p* < 0.0001, Figure 2F).

To further explore the origins of depth selectivity, we visualized running speed and optic flow speed preferences for neuronal populations preferring different virtual depths in logarithmic space (Figures 2G, S2I-J). Since different depths generate different ratios of running and optic flow speeds, log-optic flow speeds associated with each depth lie along a line as a function of log-running speed with a different intercept (Figure 2G, inset). Responses of V1 neurons mirrored this principle – neurons tuned to the same virtual depth have a wide range of preferred running and optic flow speed with a given ratio, which lie along along a line in log-running speed / log-optic flow speed space (Figure 2G). Consequently, neurons with similar depth selectively prefer different combinations of running and optic flow speeds with a fixed ratio. This may enable V1 to encode depth independent of the animal’s speed of locomotion.

### Closed loop coupling of locomotion and optic flow enables accurate representation of depth

In the experiments described above, locomotion and optic flow were coupled on fast time scales (Figure S1D). Next, we examined whether this closed loop coupling was required for conjunctive coding of optic flow and running speed underlying depth-selective responses. To break the coupling between locomotion and optic flow while still maintaining the overall statistics of the visual stimulus in open loop conditions, the movement of the mouse through the VR environment was determined by a previously recorded running trajectory instead of the current running speed (Figure 2H, S5C-D). We recorded neuronal responses during blocks of closed loop and open loop trials and compared running speed and optic flow speed preferences between these conditions. Preferred optic flow and running speeds of individual neurons were correlated between closed loop and open loop trials (Figure 2I-K, running speed *r* = 0.578, *p* < 0.0001, optic flow speed *r* = 0.700, *p* < 0.0001). The magnitude of responses at the preferred running and optic flow speeds was similar on closed loop and open loop trials (Figure 2L, *p* = 0.681). Overall these results show that the conjunctive coding of optic flow and locomotion speeds does not depend on their closed loop coupling.

Next, we asked whether the closed loop coupling of locomotion and optic flow helps maintain an accurate representation of depth at the population level. When animals run at different speeds, stimuli at different depths can give rise to the same speed of optic flow. During closed loop trials, optic flow inputs are accompanied by locomotion-related modulation corresponding to the running speed that produced them, which may help disambiguate the depth of such stimuli. On the other hand under open loop conditions, the locomotion-related modulation no longer matches the current optic flow speed. Therefore, open loop recordings allow us to experimentally “scramble” locomotion-related modulation in V1 and reveal its role in the representation of virtual depth from motion parallax. To this end, we trained a linear SVM classifier to decode virtual depth from the responses of simultaneously recorded neurons under closed or open loop conditions. Classifier accuracy was consistently higher on closed loop trials (Figure 2M-N), demonstrating that the coupling of locomotion and optic flow is necessary for accurate representation of virtual depth in V1.

### V1 neurons have three-dimensional receptive fields

We hypothesised that depth-selective neurons are driven by the presentation of visual cues at specific retinotopic locations and at specific virtual depths. To test this hypothesis, we reconstructed the stimulus presented during every imaging frame and fit neuronal responses using a linear model to estimate neuronal receptive fields (RFs) while accounting for depth (Figure 3A-B; see Methods).

**Figure 3.**
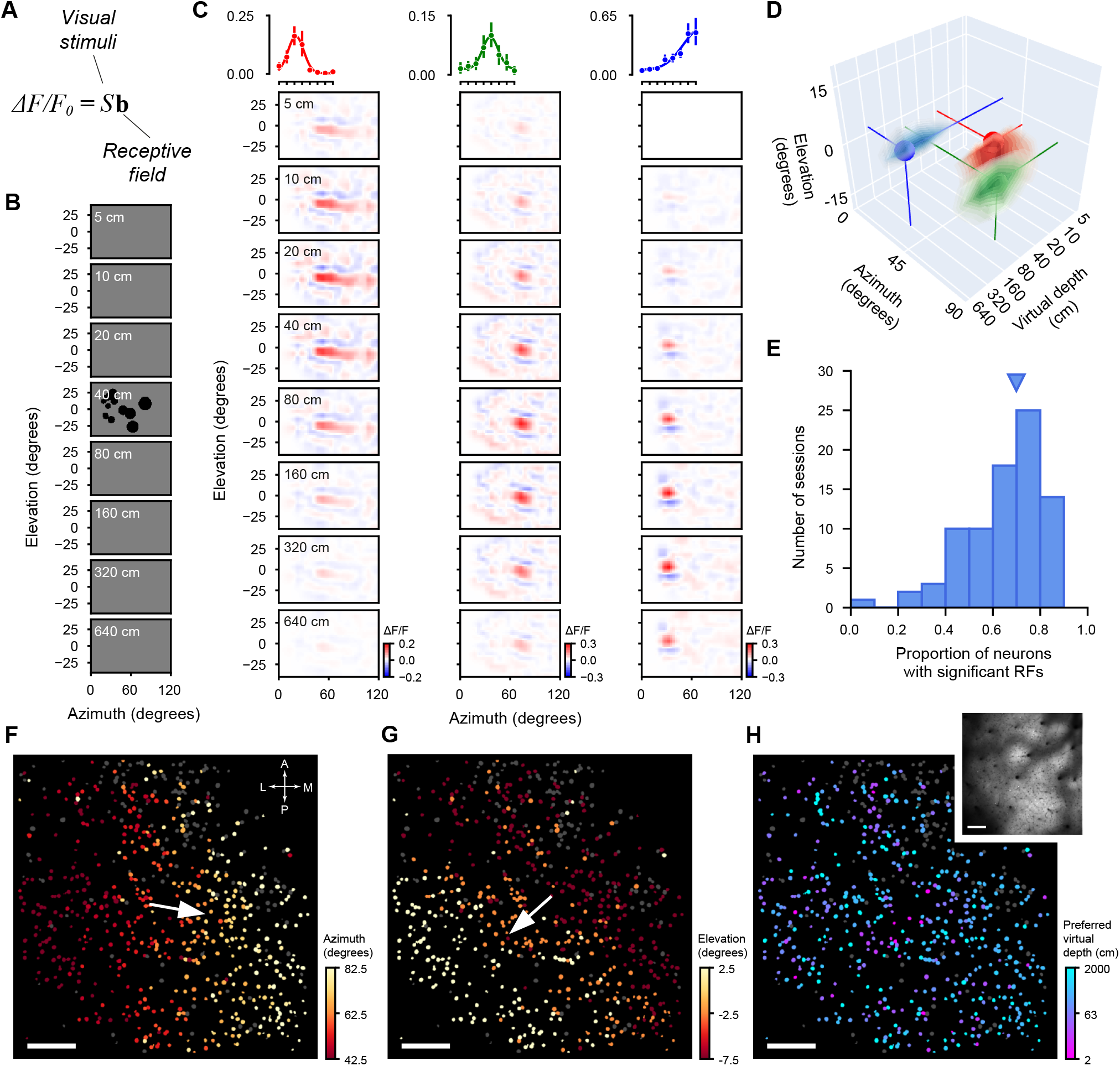
Depth-selective neurons have three-dimensional receptive fields. **(A-B)** Linear receptive field model schematic (**A**) and an example reconstructed stimulus frame (**B**). Stimuli were downsampled to 5 degrees for analysis. **(C-D)** Receptive fields of 3 example V1 neurons as a function of virtual depth **(C)** and visualized in 3D **(D)**. The rendered volume in **D** represents the receptive field locations with responses greater than half of maximum for each cell. **(E)** Proportion of neurons with significant receptive fields across imaging sessions (N = 85 sessions). Triangle indicates the median value. **(F-H)** Preferred azimuth **(F)**, elevation **(G)**, and depth **(H)** of depth-selective neurons across the field of view in an example imaging session. Scale bar – 100 *µ*m. Arrow – direction of gradient of increase in preferred azimuth or elevation. Inset – mean fluorescence image.

Many neurons responded to stimuli at specific retinotopic locations (Figure 3C). Moreover, these spatially selective responses were modulated by virtual depth in accordance with depth selectivity observed based on trial-average responses (Figure 3C). Consequently, neuronal responses could be described by a three-dimensional receptive fields (Figure 3D). Across the population, the majority (66.3%, 20,554 / 31,013) of depth-selective neurons had detectable receptive fields (Figure 3E), which represents a lower bound on the proportion of spatially tuned depth-selective neurons as receptive fields may be difficult to detect for neurons with lower signal-to-noise responses. As expected given the retinotopic organization of V1, the spatial location of RFs recorded within the same field of view was clustered in a particular region of the visual field and their preferred azimuth and elevation followed expected retinotopic gradients (Figure 3F-G, S6A-B). In contrast, depth selectivity followed a salt-and-pepper pattern, with nearby neurons often preferring different virtual depths (Figure 3H, S6C). These results demonstrate that when navigating through a three-dimensional virtual environment, responses of V1 neurons can be characterized by three-dimensional receptive fields defined by the conjunction of retinotopic location and preferred virtual depth.

### Depth selectivity across V1

We next asked how virtual depth was represented across V1. To this end, we first used widefield calcium imaging to identify the location of V1 and higher visual areas (Figure 4A). We then characterized depth-selective responses across V1 and aligned the recordings across animals to a common reference frame to visualize their distribution at different cortical locations (see Methods). Neurons’ preferred azimuth (Figure 4B) and elevation (Figure 4C) followed the expected gradients across V1. While neurons preferring different depths were intermingled, we found that neurons tuned to near virtual depths were over-represented in posterior V1 (Figure 4D). Across our dataset, the preferred virtual depth of depth-selective neurons was negatively correlated with their anterior-posterior location in V1 (*r* = − 0.239, *p* = 0.0007).

**Figure 4.**
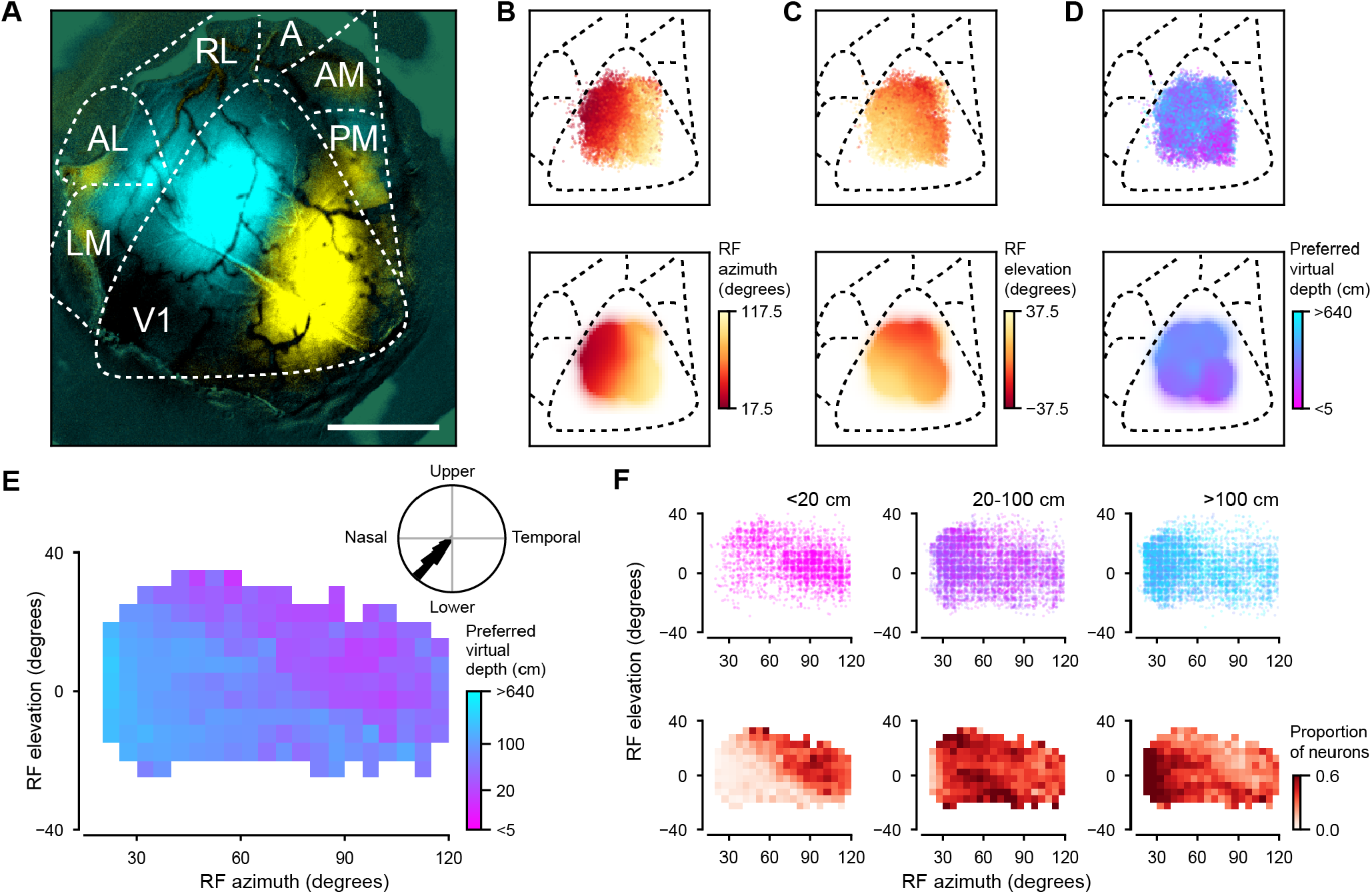
Distribution of depth preferences across the visual field. **(A)** Location of V1 and higher visual areas identified using widefield calcium imaging. Cyan and yellow indicate responses to grating patches centered at 30 and 90 degrees of azimuth. Scale bar – 1 mm. LM, lateromedial; AL, anterolateral; RL, rostrolateral; A, anterior; AM, anteromedial; PM, posteromedial. **(B-D)** Top – spatial distributions of receptive field (RF) azimuths **(B)** and elevations **(C)**, and preferred virtual depths **(D)** of depth at the location of individual neurons; bottom – RF azimuth, elevation and preferred virtual depth smoothed with a Gaussian kernel (N = 20,554 neurons from 85 sessions). **(E)** Median corrected preferred virtual depth as a function of RF location (N = 20,554 neurons from 85 sessions). Only locations with at least 20 neurons are shown. Inset – direction of the gradient of preferred virtual depth across the visual field estimated across hierarchical bootstrap samples of the dataset. **(F)** RF locations of depth-selective neurons tuned to near (<20 cm, N = 3,753 neurons), intermediate (20-100 cm, N = 8,412 neurons) and far (>100 cm, N = 8,389 neurons) virtual depths (top) after correcting for RF location, and proportion of depth-selective neurons in each category as a function of RF location (bottom). RF locations were estimated with 5 degree resolution and was plotted with 2 degree jitters.

We next asked how the preferred virtual depth of individual neurons relates to their RF location. So far, we have defined virtual depth as the virtual distance to the sphere stimuli when the animal passes them at 90 degrees of azimuth. However, the depth of the spheres varies as a function of retinotopic location as spheres that are not at 90 degrees azimuth are further away. To control for this, for each depth-selective neuron with a significant RF, we calculated the distance of the spheres at its preferred azimuth and elevation (see Methods). The resulting adjustment had the largest effect at small azimuths, where spheres were ∼ 2 times further than at the closest point at 90 degrees azimuth. However, it was substantially smaller than the >100 fold range of depths sampled across trials.

Analyzing virtual depth preferences as a function of neurons’ preferred elevation and azimuth revealed that near depths were over-represented in the upper and posterior visual field (Figure 4E). This effect was significant when controlling for variability between mice and recording sessions (*p* = 6.51 × 10^−5^, see Methods) and persisted whether or not we applied the retinotopic correction described above (Figure S6D-E, *p* = 0.00761).

While virtual depth preferences displayed a spatial bias across the visual field, neurons tuned to the same retinotopic locations displayed a broad range of preferred virtual depths, consistent with our observation that nearby neurons often have distinct depth tuning (Figures 1I, 3H, S6C). To illustrate this, we divided depth-selective neurons into 3 populations – near-(<20 cm), mid-(20-100 cm), and far (>100 cm) preferring cells. Neurons from all 3 populations were intermingled across most of the retinotopic locations sampled by our stimuli (Figure 4F). The only notable exception was the relative absence of near-preferring cells in the anterior visual field, in part as a consequence of the bias inherent to the layout of the virtual environment. Therefore, our results indicate the V1 contains a depth map of visual space from motion parallax, with most retinotopic locations represented by distinct populations of neurons tuned to near, intermediate and far depths, and over-representation of near depths in the posterior upper visual field.

## Discussion

### Representation of depth in V1

Visual responses of V1 neurons have traditionally been studied using two-dimensional visual stimuli, such as gratings or natural images. This has led to the view of V1 as a collection of spatiotemporal filters selective for specific visual features, such as spatial and temporal frequency, orientation, and direction, in the two-dimensional visual field (28–31). However, under naturalistic conditions the visual system is faced with the challenge of parsing the structure of three-dimensional visual scenes. Moreover, vision is an active sense – the optic flow generated by movements of the observer carries rich information about the organization of the surrounding space. Here we probed responses of V1 neurons in a three-dimensional virtual reality environment, where closed loop optic flow feedback based on the animal’s locomotion acted as a cue of the depth of visual stimuli. We found that a large fraction (51.4%) of V1 neurons were selectively tuned for the virtual depth of stimuli in VR. This depth selectivity was driven by motion parallax and was absent during periods when animals were stationary. Moreover, consistent with the retinotopic organization of V1, most depth-selective neurons responded to specific retinotopic locations, and could thus be characterized by three-dimensional receptive fields. Neurons with different depth preferences were intermingled, such that most retinotopic locations were represented by neurons spanning a broad range of virtual depths.

These findings support a conceptually novel view of how mouse V1 represents the visual scene in actively moving animals. They demonstrate that motion parallax generates a depth map of visual space in V1, with distinct neuronal populations responding to near versus far visual cues. We speculate that these populations may play different roles in visually guided behaviors. For example, neurons tuned to far depths may provide information about visual landmarks for navigation, while those selective for near and intermediate depths may support detection of nearby threats.

### A gradient of depth selectivity across V1

While we found that nearby neurons exhibited a broad range of virtual depth preferences, depth selectivity was not homogeneously distributed across V1. Most saliently, near depths were overrepresented in the upper lateral visual field. This contrasts to the distribution of binocular disparity preferences in the binocular region of V1, where near depths are overrepresented at low elevations (9). These differences may reflect distinct ethological roles of monocular and binocular regions in visually guided behaviors. The binocular zone appears to play a specialized role in hunting (32). During pursuit of prey, mice use their head and eye movements to maintain the location of their target in a region with minimal optic flow (20). Consequently, mice must rely on binocular cues for hunting (32) as motion parallax provides little depth information. On the other hand, the overrepresentation of near depths in the upper lateral visual field, where binocular cues are not available, may support detection of threats. Finally, overrepresentation of far depths in the lower visual field may help animals avoid hazardous falls when navigating over uneven terrain. We speculate that this bias in the distribution of near- and far-preferring neurons may reflect biases already present in the signals conveyed by the retina. Retinal ganglion cells (RGCs) are not evenly distributed across the mouse retina, with the highest density of RGCs in the ventronasal retina (33), corresponding to upper lateral visual field. This bias is shared by W3 retinal ganglion cells which are responsive to local visual motion against a featureless or stationary background (34). These cells have primarily been viewed as sensitive to object motion, such as detection of predators. However, self-generated optic flow resulting from near visual cues also gives rise to local motion distinct from background and would also likely activate these neurons.

### Function of locomotion-related signals in V1

Since its first characterization over a decade ago (21), the modulation of mouse V1 neurons by locomotion and other movements has been extensively studied (23, 35). However, its role in visual processing has remained a subject of debate (22, 24, 25, 36–38). By analyzing neuronal responses to motion parallax as a function of both running and optic flow speeds, we found that depth selective responses arise as the product of integration of these signals. Locomotion-related signals determine the gain of optic flow responses, with the largest responses occurring on coincidence of the neuron’s preferred locomotion and optic flow speeds. V1 contains a diverse population of neurons preferring a wide range of running and optic flow speeds, and the ratio between the two speeds defines their depth preferences. Therefore, we propose that locomotion-related signals in V1 support estimation of depth from motion parallax through locomotion-dependent gain modulation of optic flow responses. Gain modulation has been proposed to support coordinate transforms, such as from retinal to body-centered (39), or from to egocentric to allocentric coordinates (40).

Here, gain modulation may support the transformation from retinotopic to three-dimensional coordinates by generating set of basis functions selective for different combinations of optic flow and locomotion speeds. Downstream neurons in deep layers of V1 or in higher visual areas may generate depth-selective representations that are invariant to the animals’ speed of locomotion by integrating inputs tuned to the same depth but preferring different running speeds.

Locomotion-related modulation has been previously proposed to encode the predicted sensory feedback arising from animals’ movements (22, 41) and in combination with feed-forward visual motion signals in V1, to enable the detection of mismatches between predicted and experienced visual motion (22, 42, 43). However, predicting optic flow based on running speed requires knowledge of the depth of objects in the visual scene. Our results suggest that locomotion-related modulation enables visual cortical circuits to infer the depth of the visual cues, which could later be used to make accurate predictions of visual feedback.

### Circuit mechanisms of depth selectivity

The integration of optic flow and locomotion-related signals underlying depth-selective responses may occur in layer 2/3 of V1 *de novo* or be inherited from other sources, including the dorsal lateral geniculate nucleus (dLGN), or superior colliculus, as their spontaneous and visually-driven responses are modulated by locomotion (44–46). Several circuit mechanisms have been put forward as pathways for locomotion-related gain modulation in V1. Fu et al. proposed that locomotion modulates responses of excitatory cells through *Vip*-positive interneuron-mediated disinhibition. Leinweber et al. found that feedback projections from the secondary motor cortex are correlated with the expected optic flow resulting from animals’ locomotion, suggesting that locomotion-related modulation is conveyed by corticocortical pathways. In addition, inputs from both dLGN and the lateral posterior nucleus were shown to encode a combination of locomotion and optic flow signals (48), supporting the idea that locomotion-related modulation in depth-selective neurons may be inherited from the thalamus. Finally, locomotion has also been found to modulate effective recurrent connectivity within the cortical microcircuit (27), although the mechanism of this modulation is unclear.

Mouse primary visual cortex sends projections to several higher visual areas, which display different preferences for the speed of visual motion – anterior visual areas prefer fast moving grating stimuli, while posterior ones prefer slow ones (49, 50). As optic flow speed preferences of V1 neurons correlate with their depth selectivity, higher visual areas may be specialized for processing visual signals from different depths – with anterior areas preferentially responding to near visual cues, and posterior areas responding to far ones. Consistent with this, higher visual areas have distinct binocular disparity preferences, with anterior area RL tuned to disparities related to near visual cues (9).

### Summary

By manipulating the depth of visual cues in a virtual reality environment, we found that selectivity for depth from motion parallax is widespread in the mouse primary visual cortex. While in the virtual environment used in our experiments self-generated optic flow acts as the only source of visual motion, in the real world visual motion results from both movements of the observer and that of visual cues. Our results raise the question of how the conjunctive responses of V1 neurons enable discrimination between self-generated and external motion and support depth estimation of moving visual cues. For a moving visual stimulus, the total visual motion is the sum of external and self-generated components. When the observer varies its self-motion speed, the depth of the stimulus determines the slope of the relationship between visual motion speed and self-motion, while external motion determines the intercept. We speculate that the conjunctive code of optic flow and locomotion speed in V1 may serve as a basis set for more abstract three-dimensional representations of the visual scene further along the visual cortical hierarchy that are invariant to locomotion speed and discriminate between self-generated and external motion.

## Acknowledgments

This work was supported by the Francis Crick Institute which receives its core funding from Cancer Research UK (CC2108), the UK Medical Research Council (CC2108), and the Wellcome Trust (CC2108). We thank M. Florencia Iacaruso, Chunyu Ann Duan, and Ivana Orsolic for comments on the manuscript. We thank the Francis Crick Institute’s Making Lab, Biological Research Facility, and Scientific Computing for technical support. This preprint was typeset in LaTeX based on a template created by Ricardo Henriques.

## Author contributions

Y.H. and P.Z. designed the experiments; Y.H. and A.B. developed the experimental setup; Y.H. and A.C.N. performed the experiments; Y.H., A.B. and P.Z. analyzed the data; Y.H and P.Z. wrote the manuscript.

## Data and code availability

Preprocessed data, including fluorescence traces of individual neurons and metadata required to reproduce the analyses will be deposited on Figshare and made publicly available prior to publication. Both preprocessed and raw image data is available from Petr Znamenskiy upon request. Original code required to reproduce the analyses is available on Github at https://github.com/znamlab/v1_depth_map.

Bonsai workflows for visual stimulus presentation are available at https://github.com/znamlab/vis-stim-depth.

Original code for preprocessing of two-photon imaging data is available at https://github.com/znamlab/2p-preprocess.

## Methods

### Animals

All experimental procedures were performed in accordance to the UK Animals (Scientific Procedures) Act of 1986 (PPL PP4882546) and approved by the Animal Welfare Ethical Review Body at the Francis Crick Institute. 7 transgenic mice were used in the experiments, including 6 Emx1-Cre × Ai95D mice (JAX stock #028865 (51) and #005628 (52)) and one CamKII-tTA × TRE-GCaMP6s line G6s2 mouse (JAX stock #003010 (53) and #024742 (54)) expressing genetically encoded calcium indicators GCaMP6f or GCaMP6s respectively in cortical excitatory neurons. Both male and female mice were used (1 male and 6 females).

### Cranial window implantation

To implant the cranial window for chronic two-photon calcium imaging, mice were anesthetized with isofluorane (1.5-2.5%). They received a subcutaneous injection of dexamethasone (0.02 ml at 2 mg/ml) to reduce brain adaema. The skull was exposed and a metal headplate was affixed using dental cement. A craniotomy (4 mm in diameter) was performed over the left visual cortex and the bone flap was replaced by a 4 mm glass coverslip. Imaging was typically performed at the age of 3 to 7 months (up to 10 months).

### Virtual reality setup and visual stimuli

During the recordings, mice were head-fixed on a polystyrene cylindrical wheel and presented with a virtual reality (VR) setup on four monitor screens (MSI Optix G241) surrounding the mouse. The monitors were arranged in portrait orientation approximately along four sides of a hexagon centered on the mouse and covered a visual field of ∼ 240 degrees horizontally and ∼ 80 degrees vertically. The animals’ position in the VR environment was updated in closed loop based on the distance they traveled on the wheel recorded using a rotary encoder (Kubler 2400). The virtual reality environment was rendered in Bonsai software (55) using the BonVision package (56). The 3D environment was rendered using cube mapping, which projected a 360° visual scene onto the view ports around the mouse (56). The view ports were defined as the positions of the screens relative to the mouse, whose translation and rotational vectors were calibrated using ArUco markers (57, 58) displayed on the screens and placed at the location of the mouse. The right eye was monitored with a camera (Basler acA1440-220um) synchronized with the two-photon acquisition (∼ 15 Hz) and illuminated with an infrared light placed above the camera. The VR environment had a gray background (11.9 cd/m^2^), a 5 cm wide floor with a checkerboard texture positioned 2.5 cm below the mouse in the VR, and 24 black spheres presented at varying virtual distances from the animal. The locations of the centers of the spheres were selected in cylindrical coordinates (Figure S1A). The longitudinal axis of the cylinder was aligned to the animals path of travel in the VR, while its radius was equal to the virtual depth of a given trial chosen pseudorandomly from a range of logarithmically spaced values (either 5 virtual depth values between 6 cm and 600 cm, or 8 depths between 5 and 640 cm). The spheres were spaced along the axis of the animal’s locomotion by 0.15 × *d* with 19 spheres ahead of and 5 spheres behind the mouse. The cylindrical azimuth of the spheres, which determines their position relative to the horizon, was randomly chosen from a uniform distribution between -40 and 40 degrees or between 140 and 220 degrees, i.e. around the horizon on either the left-hand side or right-hand side of the animal (Figure S1A). To maintain constant visual stimulation during the trial, whenever the animal moved the distance of 0.15 × *d* in the VR, resulting in a sphere disappearing outside of the monitors’ field of view behind the mouse, a new sphere was generated 2.7 × *d* ahead of the mouse at a randomized cylindrical azimuth. New spheres gradually faded in from gray over 0.3 s. The virtual radius of the spheres on trials of different virtual depths was equal to tan 5° × *d*, so that they covered the same angular extent in the visual field of the mice across different virtual depths (10 degrees when the mice passed closest to each sphere in the VR). The spheres were a solid black color without specular highlights.

Each trial continued until the mouse traveled 6 m after which the spheres faded to gray and a 10 s inter-trial period commenced. A probabilistic reward of soy milk (∼ 8 *µ*l of 10% SMA Wysoy) was delivered from a spout in front of the mouse 2 s into the inter-trial interval on 60-80% of trials. Each session consisted of at least 10 trials for each virtual depth.

During open loop sessions, the animal’s movement on the wheel was recorded but did not control its virtual position in VR. Instead, the running trajectory of a previous closed loop session of the animal was replayed and determined the animal’s moment-to-moment virtual position and the updating of the sphere stimulus.

The nominal refresh rate of the four monitors was 144 Hz. To synchronize the visual stimuli with imaging data, and measure the actual frame rate, a small square (3 cm × 3 cm) with varying luminance was displayed in the bottom left corner of the leftmost monitor (Figure S1B). The luminance changes of this square were recorded with a photodiode at 1 kHz (HARP Behavior device, Champalimaud Research Scientific Hardware Platform). For 2 mice (25 sessions), the square alternated at every frame between black and white and was used to detect dropped frames. 89.7 *±* 5.1% of the frame were presented at 144 Hz, resulting in a average frame rate per session above 124 *±* 9 Hz. To determine the closed loop latency between rotary encoder inputs and updates of the visual stimulus, for the remaining 5 mice (60 sessions), we updated the brightness of the square following an irregular predefined sequence of 5 values (Figure S1B). The sequence was selected to ensure that brightness increases and decreases were alternating at each frame. We then detected frames on the photodiode trace and computed the cross-correlation between the filtered predefined sequence and the photodiode recording to identify which frame was presented at each time point and to continuously measure display lag. The average display lag per session was 26.6 ± 1.4 ms (Figure S1C-D).

### Behavioral training

To encourage mice to run in the VR environment, they underwent a 2-4 week training period under food restriction. Mice were acclimatized to the behavioral setup for at least 3 sessions of 5-15 min. Mice initiated their training with a minimum distance traveled per trial of 1 m that was gradually increased to 6 m depending on the animals’ running behavior. Each training session lasted 30 min - 1 h. Imaging commenced once the mice could reliably complete ∼ 2 trials per minute. After the training period, food restriction continued during the imaging experiments.

### Two-photon calcium imaging

Fluorescence signals from neurons in layer 2/3 of the primary visual cortex (150-300 *µ*m below pia) were recorded using a custom-built resonant scanning two-photon microscope (Independent NeuroScience Services) and a Nikon 16x water-immersion objective (NA 0.8). The microscope was controlled with ScanImage 2020 software. GCaMP6f and GCaMP6s fluorescence signals were excited with a 930 nm laser (SpectraPhysics MaiTai DeepSee eHP) at 70-100 mW and acquired with a 525/50 emission filter (FF03-525/50-25, Semrock). Imaging frames (1024 × 1024 pixels) were acquired at ∼15 Hz. Individual recordings covered a field of view of ∼ 661 *µ*m × 661 *µ*m. To minimize the interference from the monitors’ during imaging, a custom electronic circuit activated the backlights only during the turn-around period of the resonant X mirror.

### Widefield calcium imaging

The locations of V1 and higher visual areas (HVAs) were determined using widefield calcium imaging of the whole visual cortical window. Widefield calcium imaging was performed using a camera integrated into the two-photon microscope (Independent NeuroScience Services) using a Nikon AF Nikkor 85mm f/1.8D camera lens as the objective and a 100 mm tube lens (Thorlabs AC508-100-A-ML). Two excitation LEDs at 470 nm (M470L3, Thorlabs, with excitation filter FF02-447/60-25, Semrock) and 405 nm (M405L4, Thorlabs, with excitation filter FF01-405/10-25, Semrock) were combined using a dichroic mirror (FF458-Di02-25×36, Semrock) and coupled to the infinity focused imaging path (FF495-Di03 BrightLine primary dichroic mirror). Images were acquired with an emission filter (FF03-525/50-25, Semrock) and a CMOS camera (Basler acA2040-120um) at 60 Hz. Illumination of the two LEDs was interleaved for hemodynamic correction using a microcontroller (Arduino Uno) triggered by the camera’s FlashWindow output.

During imaging, mice were head-fixed on the wheel and presented with a circular square-wave grating patch of 20 degrees diameter (at a spatial frequency of 0.2 cpd and a temporal frequency of 2 Hz) at 30 or 90 degrees azimuth and 0 degrees elevation on a black background. Each grating patch was presented for 4 seconds and grating direction (8 directions in 45 degree increments) changed every 500 ms in pseudorandom order. The grating patches were repeated 50 times for each location in pseudorandom order with 8 seconds of blank screen between presentations.

Images were first deinterleaved to separate 405 nm and 470 nm channels. Hemodynamic signal correction was carried out by performing linear regression on each pixel of the 470 nm channel’s signal using the 405 nm channel signal as the independent variable. The residuals of the regression added to the mean fitted signal of the 470 nm channel was used as the corrected widefield signal for each pixel. Corrected signals were detrended by applying a high-pass filter of 0.001 Hz. The signal at each pixel was normalized by its average value. The responses to the grating patches at each retinotopic location were averaged across repetitions to determine the locations of V1 and HVAs.

## Data analysis

To account for the nested structure of the experimental data, statistical comparisons were conducted using hierarchical bootstrap (59) unless specified otherwise. In analyses at the level of individual neurons, for each bootstrap sample we resampled mice, followed by experimental sessions for each mouse, and neurons for each session and calculated the statistic of interest, such as correlation coefficient between two variables, or difference in medians between two conditions, for each sample. We then determined the quantile *q* of the bootstrap distribution at 0 and calculated the two-sided bootstrap p-value as 2 min{*q*, 1 −*q*}.

### Eye movements

To determine if the visual stimulation triggered eye movements, in a subset of sessions (N = 16 sessions from 2 mice) we reconstructed the gaze direction during behaviour. Eyelid position, the reflection of the infrared light and 12 key points around the pupil border were tracked on each frame using DeepLabCut (60). Frames in which the eye was closed (distance between top eyelid and bottom eyelid < mean distance - 3 standard deviations, or DeepLabCut likelihood <0.88) were excluded. For the remaining frames, the 12 markers around the pupil were fitted with an ellipse which was used to perform gaze reconstruction as previously described (61). Briefly, the eye was modeled as a 3D sphere with a circular pupil rotating at a constant radius around the the eye center. Translation of the eye relative to the camera was corrected using the infrared light reflection as the origin for each frame. The position of the eye center on the camera frame was defined as the least square estimate of the intersection of all minor axes of the fitted ellipses. The scale factor, which depends on the radius of the eye, was estimated based on the ratio of the short and long axis of the fitted ellipses and the position of the eye center (61). With these parameters, the pupil radius *r* and rotation angles relative to the camera axes, *ϕ* and *θ* define the reprojected pupil border in the camera frame. We performed a grid search for each frame to find the (*r, ϕ, θ*) minimizing the difference between the fitted and reprojected ellipses.

To estimate the azimuth and elevation from the gaze vector in camera coordinates, we calibrated the camera position using an ArUco marker (58) placed parallel to the floor in the camera field of view and pointing in the animal’s direction of travel. The pose of the marker relative to the camera was estimated using OpenCV (57) and used to rotate the gaze vector from camera coordinates to world coordinates.

Angular velocity (Figure S2K) was calculated on the mean filtered azimuth and elevation traces (rolling window of 3 frames). Saccades (Figure S2L) were defined as instances when the filtered velocity traces (calculated on median-filtered azimuth and elevation traces with a rolling window of 5 frames) exceeded 75 degrees per second, i.e. a 5 degrees change in a frame.

### Preprocessing of two-photon imaging data

Imaging frames were registered and segmented using the suite2p package (62) with anatomical segmentation (Cellpose (63)) to avoid bias towards active cells. Fluorescence traces were detrended by subtracting a rolling baseline calculated as the 20th percentile in a moving window of 900 frames. To remove neuropil contamination of the fluorescence trace for the ROIs, we fitted the fluorescence of the ROIs and their surrounding areas with asymmetric Student-t distributions. The mean of these distributions depended on a shared neuropil signal that affected both ROI and surrounding fluorescence (64). Finally, we estimated *F*_0_, by fitting a Gaussian mixture model with 2 components to the fluorescence time series and identified *F*_0_ as the mean of the smaller Gaussian. Δ*F* /*F*_0_ was defined as Δ*F/F*_0_ = (*F* ™ *F*_0_)*/F*_0_, where *F* was the fluorescence value after neuropil correction.

### Depth selectivity

To determine the preferred virtual depth for each ROI, we calculated the mean Δ*F* /*F*_0_ across each trial for each neuron, and fitted a Gaussian model to the average single trial responses *f* and the corresponding log-transformed virtual depth *d* displayed in each trial:

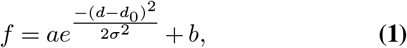

where *a* was the peak response amplitude, *d*_0_ was the log-preferred depth, *σ* was the tuning width, and *b* was the baseline fluorescence. The preferred virtual depth *d*_0_ was constrained between ln 2 and ln 2000 cm, and the peak amplitude *a* was constrained to be positive. The tuning width *σ*^2^ was constrained to >0.5 to avoid overfitting. All depth tuning curves were plotted using this fit.

To identify depth-selective neurons, we fitted the Gaussian model using 5-fold cross validation. On each fold, 80% of trials were assigned to the training set, which was used to estimate model parameters and 20% were assigned to the test set, which was used to evaluate model predictions. Then we calculated the Spearman’s correlation coefficient of predicted and observed fluorescence on test trials across all 5 folds. Depth-selective neurons were defined as having a Spearman’s correlation p value <0.05 and the Spearman’s correlation coefficient >0.1.

Near-preferring cells and far-preferring cells (Figure 1K) were defined as the cells with preferred virtual depths close to the bounds of *d*_0_ (log-preferred virtual depth < ln 2 + 10^−4^ for near-preferring cells, and *>* ln 2000 − 10^−4^ for far-preferring cells).

### Optic flow and running speed tuning

To visualize the optic flow speed and running speed tuning curves in Figure 2B and S4B,F, we first smoothed the sum of responses from all frames and the number of frames at different bins of running speed or log-transformed optic flow speed respectively, using a 1D Gaussian kernel with a standard deviation of 1 bin width (10 cm/s for running speed, ∼ 0.691 for log-transformed optic flow speed). The tuning curves were calculated by dividing the smoothed sum of response by the smoothed number of frames to account for the number of samples in different bins. Only frames with a running speed above 1 cm/s were included for the optic flow speed tuning curves to facilitate the logarithmic transformation of the optic flow speeds.

To determine how optic flow and locomotion-related signals were integrated and estimate preferred optic flow and running speeds, we fitted the Δ*F/F*_0_ trace as a function of log-transformed optic flow speed *v* and running speed *r* on individual imaging frames when mice were moving at >1 cm/s using 5 models.

The optic flow only and running speed only models were Gaussian functions of log-optic flow speed *v* and log-running speed *r*, respectively:

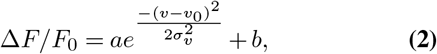

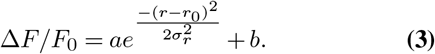

The additive model fitted responses as the sum of Gaussian functions of log-optic flow speed *v* and log-running speed *r*:

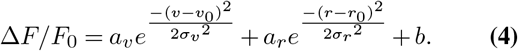

The conjunctive model consisted of a bivariate Gaussian tuned to both log-optic flow speed *v* and log-running speed *r*:

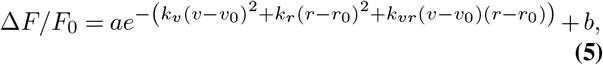

Where

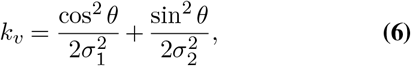

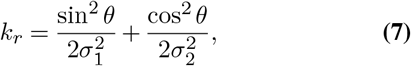

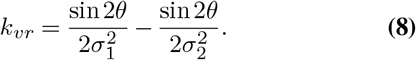

*a, a*_*r*_, and *a*_*v*_ were the peak amplitude of the overall response and the running speed and optic flow components, respectively. *σ*_*v*_ and *σ*_*r*_ were the tuning width for optic flow and running speed respectively, *b* was the baseline activity, and *v*_0_ and *r*_0_ were the log-preferred optic flow and log-preferred running speeds. For the conjunctive model, *θ* determined the orientation of the axes of the bivariate Gaussian, and *σ*_1_ and *σ*_2_ were the standard deviation for its two axes.

The ratio model was a Gaussian function of the difference between log-running speed and log-optic flow speed, equivalent to the logarithm of their ratio:

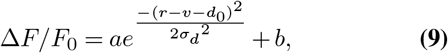

where *d*_0_ was the log-preferred virtual depth and *σ*_*d*_ was the depth tuning width. This model is similar to the depth tuning model in Eq. 1 fitted on individual imaging frames rather than trial-averages.

The preferred optic flow speed 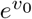 was constrained between 0.03 to 3000 degrees/s. The preferred running speed 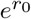 was constrained between 0.5 to 500 cm/s. *d*_0_ was constrained between 0.0095 and 9.5 × 10^5^ cm according to the lower and upper bounds of 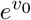 and 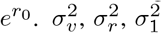, and 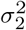 were constrained to be >0.25. *θ* was constrained between 0 and 90 degrees. *a, a*_*r*_, and *a*_*v*_ were constrained to be >0.

To facilitate the comparison of the models in Figure 2E, we selected strongly responsive neurons defined as depth-selective neurons (defined as above) with a peak response at their preferred depth >0.2 (6,277 of 31,013 depth-selective neurons from 85 sessions). To determine which model best described neuronal activity for individual neurons, we fitted each model with 5-fold cross validation. On each fold, individual trials were assigned to the training set (80%), which was used to estimate model parameters, or test set (20%), which was used to evaluate model predictions. We then computed the predicted fluorescence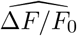 on test trials using parameters estimated on the training set for each fold and compared it to observed Δ*F/F*_0_ to compute *R*^2^ as the fraction of variance explained by the model:

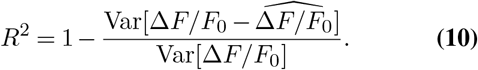

The best model for each neuron was selected as the model with the highest *R*^2^. To compare performance between models across the dataset, we first calculated the proportion of neurons best fitted by each model for each recording session and then used hierarchical bootstrap (59) to resample mice and sessions 20000 times. We then compared the difference of the median proportions between models.

To estimate the preferred running speed and optic flow speed for each neuron, we fitted the Δ*F* /*F*_0_ trace, the corresponding optic flow speed and running speed from all trials using the conjunctive model. Closed loop recordings and open loop recordings were fitted separately. The neurons plotted in Figure 2F-G,J-L were the depth-selective neurons well-fitted by the conjunctive model in closed loop recordings (cross-validated *R*^2^>0.02 computed as described above, 5,737 of 31,013 depth-selective neurons recorded in closed loop conditions from 84 out of 85 sessions; no neurons passed this threshold in 1 session). To quantify the correlation between preferred depth, preferred running speed, preferred optic flow speed or the ratio between preferred running and optic flow speed in Figure 2F and Figure S4I, J, we first calculated the Spearman’s correlation coefficient between the two variables and calculated the p-value using hierarchical bootstrap by resampling mice, sessions, and neurons as described above.

To compare the preferred running speed and preferred optic flow speed of depth-selective neurons in closed loop vs. open loop trials in Figure 2J-K, we chose depth-selective neurons with good fits to the conjunctive model (cross-validated *R*^2^>0.02) in both closed and open loop conditions (1,234 of 10,850 depth-selective neurons recorded in open loop from 34 sessions). Then, we computed the Spearman’s correlation coefficient between the variables and calculated the p-value using hierarchical bootstrap by resampling mice, sessions, and neurons as described above.

To compare the peak response amplitudes of depth-selective neurons in closed loop vs. open loop trials in Figure 2L, we calculated the peak response amplitude using the conjunctive model fit (sum of amplitude *a* and offset *b*). The selection criteria for neurons was the same as Figure 2J-K. Then we compared the median ratio between the peak response amplitude of closed loop and open loop trials to 1 using hierarchical bootstrap by resampling mice, sessions, and neurons as described above.

### Receptive field estimation

To estimate the receptive field for each neuron, we used the recorded sphere locations and the animal’s position in the VR to reconstruct the three-dimensional visual stimulus presented for each imaging frame. The first two dimensions corresponded to retinotopic location in spherical coordinates with a resolution of 5 degrees in azimuth and elevation, while the third dimension corresponded to the virtual depth of the stimulus on a given trial. We then constructed a design matrix representing the visual stimuli *S*, where each row contained the stimulus reconstruction for the corresponding imaging frame and for each virtual depth, as well as a column of 1s to account for a bias term. The number of rows corresponded to the number of imaging frames. Therefore, the dimension of *S* was *N*_frames_ ×(*N*_pixels_ * *N*_depths_) + 1, where *N*_pixels_ was the number of pixels on each visual stimulus frame, *N*_depths_ was the number of virtual depths, and the *N*_frames_ was the number of imaging frames. *S* was reconstructed based on the stimuli presented on either the two screens on the right-hand side of the mouse or the two screens on the left-hand side of the mouse. We used a linear model to fit Δ*F/F*_0_:

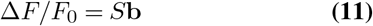

We used the regularized pseudoinverse method (65, 66) to impose a smoothness constraint on the receptive field in azimuth and elevation and in virtual depth. The constraints aimed to make the Laplacian of the receptive field close to zero at all points. To impose this constraint, we constructed matrices *L*_*xy*_ and *L*_*depth*_. *L*_*xy*_ aimed to smooth the receptive fields across azimuth and elevation. To assemble the matrix *L*_*xy*_, we first generated Laplace matrices that contained the values

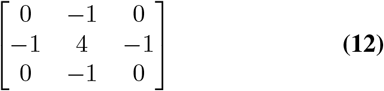

corresponding to adjacent *x* and *y* pixels for a single depth embedded in *N*_*y*_ × *N*_*x*_ × *N*_depth_ matrix of zeros for each *x* and *y* pixel location and depth. These matrices were flattened and assigned to individual rows of *L*_*xy*_.

*L*_*depth*_ controlled the smoothness the receptive fields across different virtual depths. To assemble *L*_*depth*_, we generated Laplace matrices that contained the values [−1 2 −1] corresponding to adjacent depths for a single pixel embedded in a *N*_*y*_ × *N*_*x*_ ×*N*_depth_ matrix of zeros for each *x* and *y* pixel location and depth. These matrices were flattened and assigned to individual rows of *L*_*depth*_.

We appended a column of 0s to *L*_*xy*_ and *L*_*depth*_ so as not to regularize the bias term. We then assembled the design matrix *X* containing the stimulus matrix *S* and the smoothness constraints *L*_*xy*_ and *L*_*depth*_:

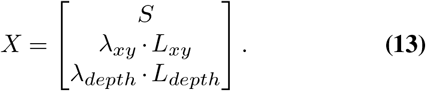

*λ*_*xy*_ and *λ*_*depth*_ are scalars that control the strength of regularization of receptive field smoothness across azimuth and elevation (*λ*_*xy*_) and across virtual depths (*λ*_*depth*_). We constructed an augmented response vector by appending 0s to the Δ*F/F*_0_ fluorescence vector corresponding to each row of *L*_*xy*_ and *L*_*depth*_

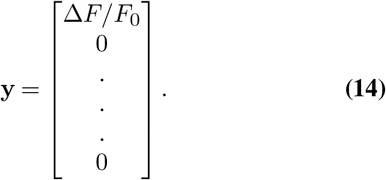

To estimate the receptive field we found the least squares solution for the equation **y** = *X***b** as

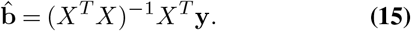

The first *N*_pixels_ elements of 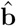 correspond to coefficients of the receptive field, while the last element represents the bias term.

To optimize hyperparameters *λ*_*xy*_ and *λ*_*depth*_, we used 5-fold cross validation to split imaging trials into training and test sets and estimated 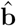 on each training set. We then computed the predicted fluorescence

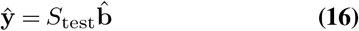

for each test set and found the combination of *λ*_*xy*_ and *λ*_*depth*_ which yielded the highest fraction of variance explained by the model for each ROI by searching over a 13 × 13 grid of values logarithmically spaced between 2.5 and 10240. The receptive field for each neuron was calculated based on the best combination of *λ*_*xy*_ and *λ*_*depth*_ by averaging the coefficients 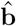 across 5 folds. The peak azimuth and elevation of the receptive field (RF_azi_ and RF_ele_) were defined as the position with the maximum coefficient value.

To identify the neurons with significant receptive fields, we used the procedure above to optimize *λ*_*xy*_ and *λ*_*depth*_ and fit the receptive field using the stimuli in the right visual field (RF_right_). To estimate the null distribution for receptive field coefficients for each neuron, we used these values of *λ*_*xy*_ and *λ*_*depth*_ to fit the receptive field using stimuli in the left visual field (RF_left_), which have the same statistics but are not expected to drive neuronal activity recorded in the left visual cortex. We calculated the mean and standard deviation of the coefficients in RF_left_ and defined the ROIs with significant receptive fields as those with a maximum value of the RF_right_ which was 6 standard deviations above the mean of RF_left_. The 6 standard deviations threshold was chosen such that ∼ 5% of neurons passed this threshold if it was applied to RF_left_.

### Cortical location

To determine the spatial location of imaged neurons within V1 in individual animals, we first found the location of neurons within the field of view (FOV) for each recording session based on their cell masks. We then aligned the location of each FOV within the whole visual cortical window by matching the blood vessel patterns between the surface of the FOVs and the overview image of the window acquired using the widefield camera.

To align the location of imaged neurons across animals, we assumed that the RF azimuth and elevation followed the same linear gradient in V1 across animals. Under this assumption, the overview location *x*_overview_ and *y*_overview_ of individual neurons depends linearly on their RF azimuth and elevation with a translational offset that can vary across mice:

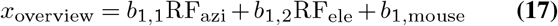

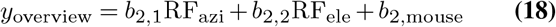

We used Huber regression to fit the coefficients and individual mouse offsets ensuring robustness to outliers. We then chose a reference mouse and aligned the neurons’ coordinates of all other mice to the reference mouse by subtracting the offset of the original mouse and adding back the offset of the reference mouse:

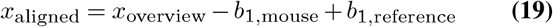

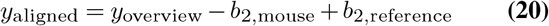

### Depth selectivity in relation to RF location

To illustrate the spatial distribution of RF azimuth, elevation and the preferred depth in Figure 4B-D, we plotted these parameters of all depth-selective neurons with a significant RF at the location of individual neurons. In the lower panels, we smoothed the spatial distribution map with a 113 *µ*m Gaussian kernel. To set the transparency of the smoothed map, we calculated the sum of Gaussian weights for all plotted neurons for each pixel, normalized by its maximum value. The alpha for each pixel was set to either 5 times this value or 1, whichever was smaller.

As the virtual distance between the spheres and the mouse varied based on the spheres’ positions, in the analyses in in Figure 4E-F we calculated the corrected log-preferred virtual depth 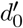, corresponding to the distance to the spheres at the center of the neurons’ receptive field at the peak of their depth tuning curve (Eq. 1):

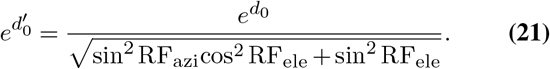

To test for the presence of a gradient in preferred depth as a function of azimuth and elevation, we used hierarchical bootstrap (59) to resample neurons across mice and recording sessions. For each bootstrap sample, we performed linear regression between the RF_azi_ and RF_ele_ of the center of the neurons’ receptive fields and the neurons’ corrected log-preferred virtual depth:

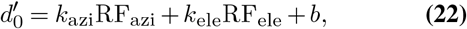

where *k*_*azi*_ and *k*_*ele*_ defined the gradient of depth as a function of azimuth and elevation of the neurons’ receptive fields, respectively.

To test for statistical significance, we modeled the coefficients (*k*_azi_, *k*_ele_) obtained from different bootstrap samples using a bivariate Normal distribution and used a multivariate generalization of the Z-test to compare their mean to 0. To this end, we first computed the mean ***µ*** and covariance Σ of bootstrap samples (*k*_azi_, *k*_ele_). As the square of the Mahalanobis distance ***µ***^*T*^ Σ^−1^***µ*** follows the chi-squared distribution with 2 degrees of freedom, the p-value is calculated as 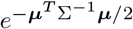

### Population decoding

To decode virtual depth from population activity, we trained a linear SVM classifier using Δ*F* /*F*_0_ values of simultaneously recorded neurons in a session. We split imaging trials into training (64%), validation (16%) and test (20%) sets with 5-fold cross validation such that each trial was included in one test set. We then trained the classifier on individual imaging frames from trials in the training set using the validation set to optimize the hyperparameter *C*. We used the test sets to evaluate the performance of the optimized classifier to obtain the accuracy and the confusion matrix of predicted and true virtual depths. Confusion matrices in Figures 1M and 2N show decoder performance as the proportion of imaging frames for each true depth.

To determine how decoding accuracy depends on the animals’ running speed, we trained a single decoder using data across all running speeds for each session. We then evaluated decoder performance on imaging frames from trials in the test set belonging to different running speed bins. Figure 1M shows mean decoding error across sessions. Confidence intervals were computed using bootstrap by resampling sessions.

To compare decoding accuracy between closed loop and open loop trials, we trained and evaluated separate SVM classifiers on closed loop and open loop data.

**Figure S1.**
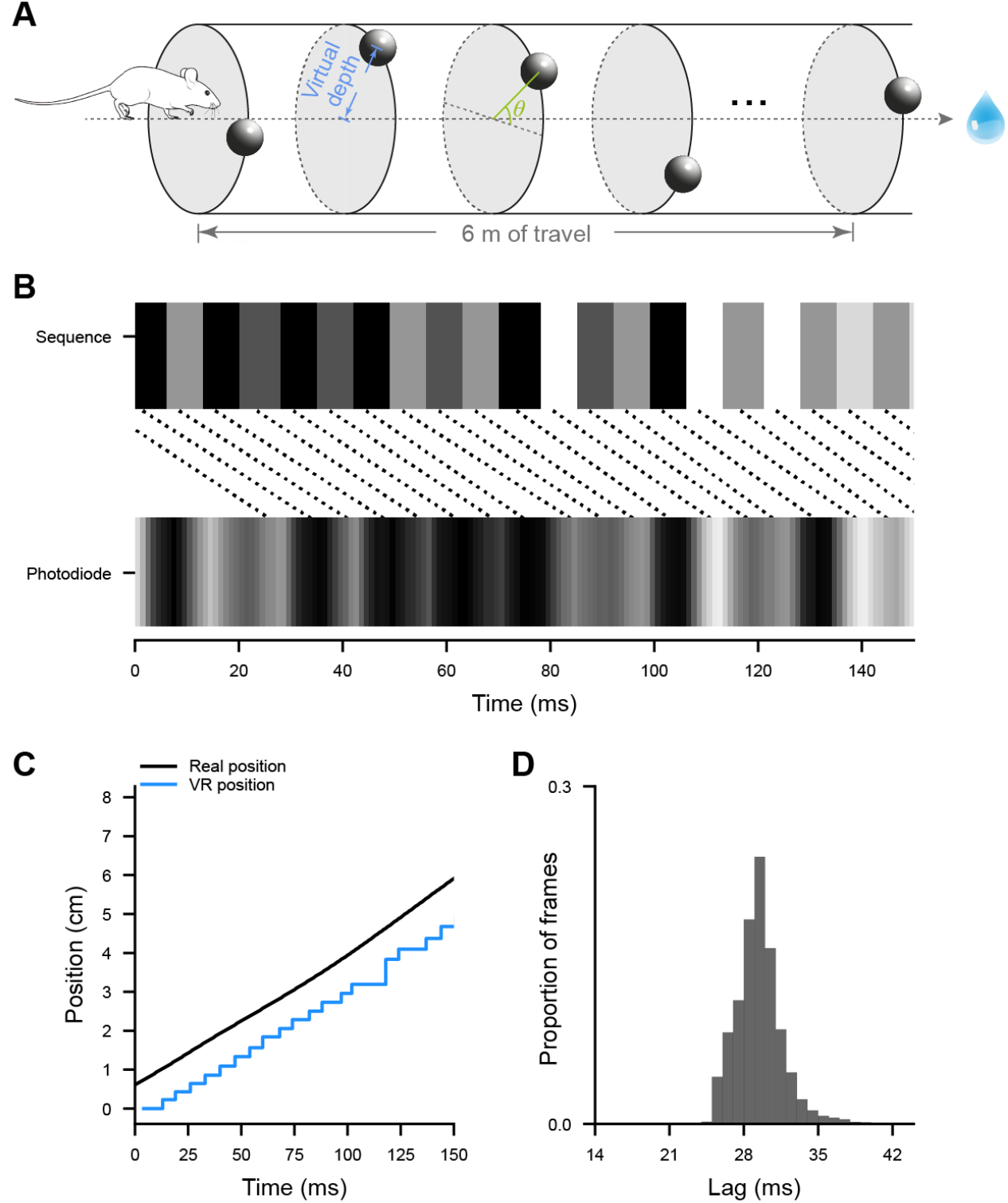
Visual stimuli in virtual reality. **(A)** Schematic illustrating the generation of sphere locations. The locations of the centers of the spheres were selected in cylindrical coordinates. Virtual depth determined the radius of the cylinder, while cylindrical azimuth of the spheres *θ* (selected from a uniform distribution between -40 and 40 degrees or between 140 and 220 degrees) determined their position relative to the horizon. The mouse drawing from Tyler, E., & Kravitz, L. (2020), Zenodo.doi.org/10.5281/zenodo.3925915. **(B)** Example sequence of varying luminance presented in the corner of the monitor for synchronization (top) and the recorded photodiode signal (bottom). Dashed line indicates the corresponding frames. **(C)** Real time mouse position (black), as measured by the rotary encoder, and the current mouse position used to render the VR environment (blue). Each step on the blue trace is a frame and the horizontal shift between the black and blue traces is the closed loop latency. **(D)** Histogram of VR latencies at each frame.

**Figure S2.**
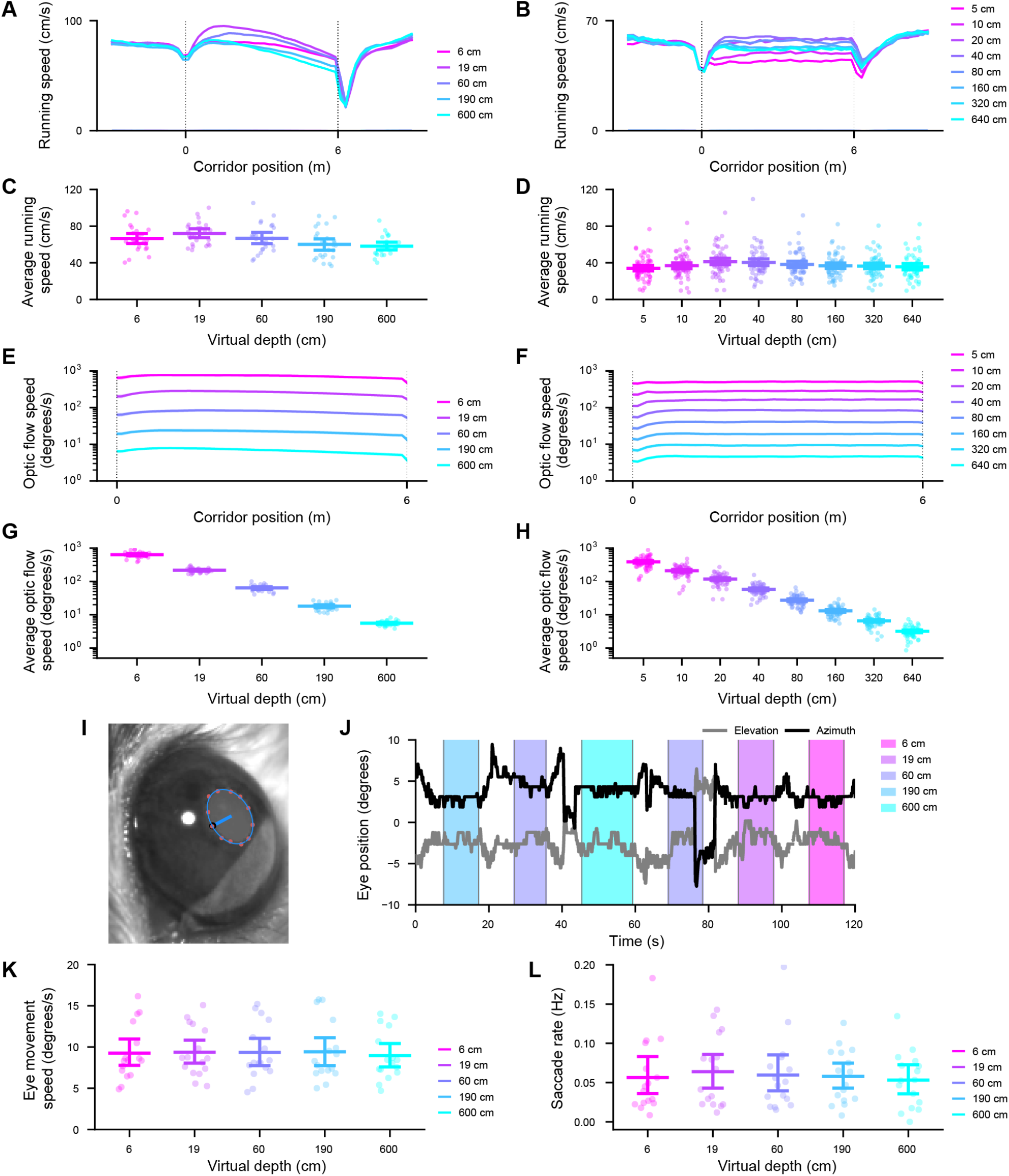
Running, optic flow speeds and eye movements across different virtual depths. **(A-B)** Mean of session average running speed as a function of distance traveled during the trials at 5 (**A**, N = 25 sessions) or 8 (**B**, N = 60 sessions) virtual depths, including 3 m of inter-stimulus interval before and after the trials. Shading – 95% confidence interval. **(C-D)** Mean running speed of each session at 5 **(C)** or 8 **(D)** virtual depths. Error bar – 95% confidence interval. **(E-F)** Mean of session average optic flow speed as a function of distance traveled during the trials at 5 (**E**) or 8 (**F**) virtual depths. Shading – 95% confidence interval. **(G-H)** Mean optic flow speed of each session at 5 **(G)** or 8 **(H)** virtual depths. Error bar – 95% confidence interval. **(I)** Example frame of the right eye recorded during imaging. The pupil border was tracked using DeepLabCut (orange dot), fitted with an ellipse (blue circle) and gaze (blue line) from the eye center (black dot) was estimated as described in (61). **(J)** Example gaze direction in azimuth and elevation relative to median direction during a recording. **(K-L)** Average eye velocity **(K)** and saccade rate **(L)** by session across trials of different virtual depths (N = 16 sessions, 2 mice).

**Figure S3.**
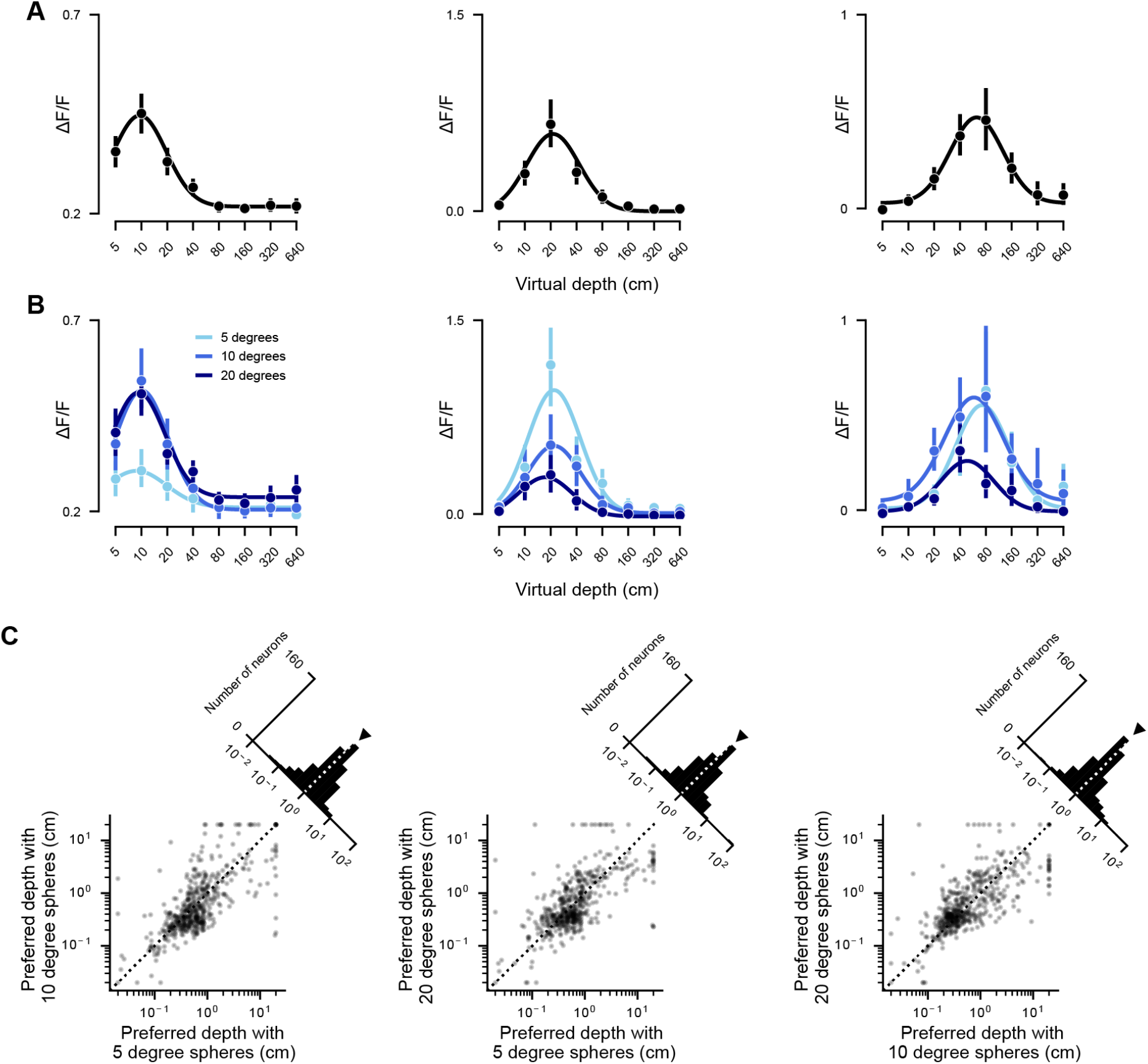
Virtual depth selectivity is invariant to stimulus size. **(A)** Virtual depth tuning of 3 example neurons across all stimulus sizes. **(B)** Virtual depth tuning of the neurons in (A) in response to different sizes of sphere stimuli. **(C)** Preferred virtual depths of all depth-selective neurons (500 neurons from 3 sessions in 2 mice) across pairs of stimuli sizes. Triangles indicate medians. Insets – histograms of the ratio of preferred virtual depths across pairs of stimuli sizes. Preferred virtual depth mapped with 5 degree vs. 10 degree spheres, median ratio = 1.13, *p*_*ratio*_ = 0.232, *r* = 0.678, *p*_*correlation*_ < 0.0001; 5 vs. 20 degree spheres, median ratio = 1.13, *p*_*ratio*_ = 0.401, *r* = 0.635, *p*_*correlation*_ < 0.0001; 10 vs. 20 degree spheres, median ratio = 0.93, *p*_*ratio*_ = 0.229, *r* = 0.721, *p*_*correlation*_ < 0.0001.

**Figure S4.**
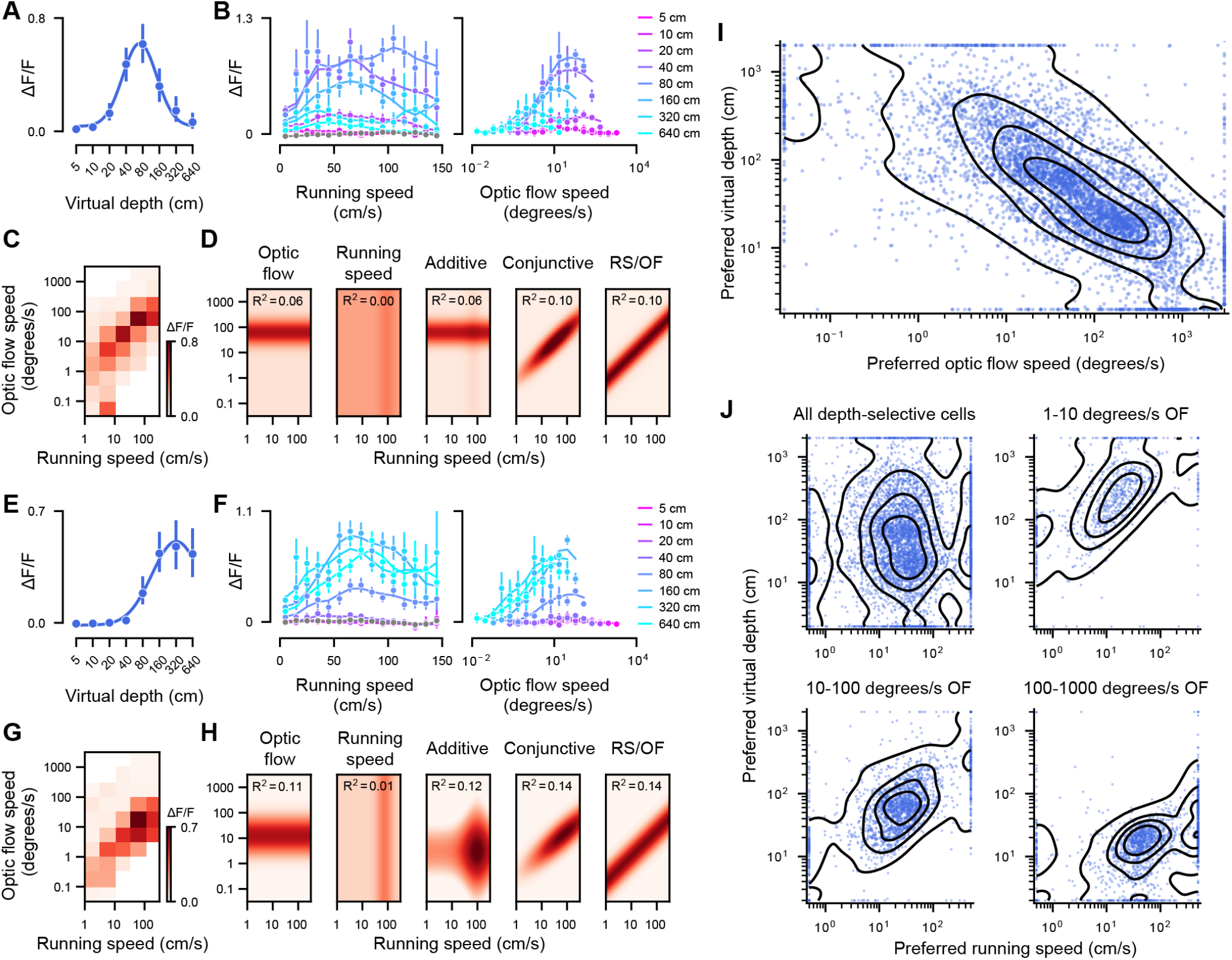
Depth selectivity arises from a conjunctive coding of optic flow speed and running speed. **(A-B)** Virtual depth tuning **(A)** and responses of an additional example depth-selective neuron binned by running speed (**B** left) or optic flow speed(**B** right). Gray line (**B** left) – running speed tuning during the inter-trial interval, when sphere stimuli were absent. **(C)** Responses of the example neuron in **A-B** and as a function of both running and optic flow speeds. **(D)** Running speed and optic flow tuning fitted with the four models of the example neuron in **A-C**. RS – running speed, OF – optic flow speed. **(E-H)** Responses of an additional example neuron as in **(A-D). (I)** Preferred virtual depth as a function of preferred optic flow speeds estimated using hold-out data not used in calculating preferred virtual depths (*r* = −0.728, *p* < 0.0001). **(J)** Preferred virtual depth as a function of preferred running speeds of all neurons estimated using hold-out data not used in calculating preferred virtual depths (top left, *r* = 0.013, *p* = 0.715), and for neurons with preferred optic flow speeds between 1–10 degrees/s (top right, *r* = 0.617, *p* < 0.0001), 10–100 degrees/s (bottom left, *r* = 0.568, *p* < 0.0001), and 100–1000 degrees/s (bottom right, *r* = 0.489, *p* < 0.0001).

**Figure S5.**
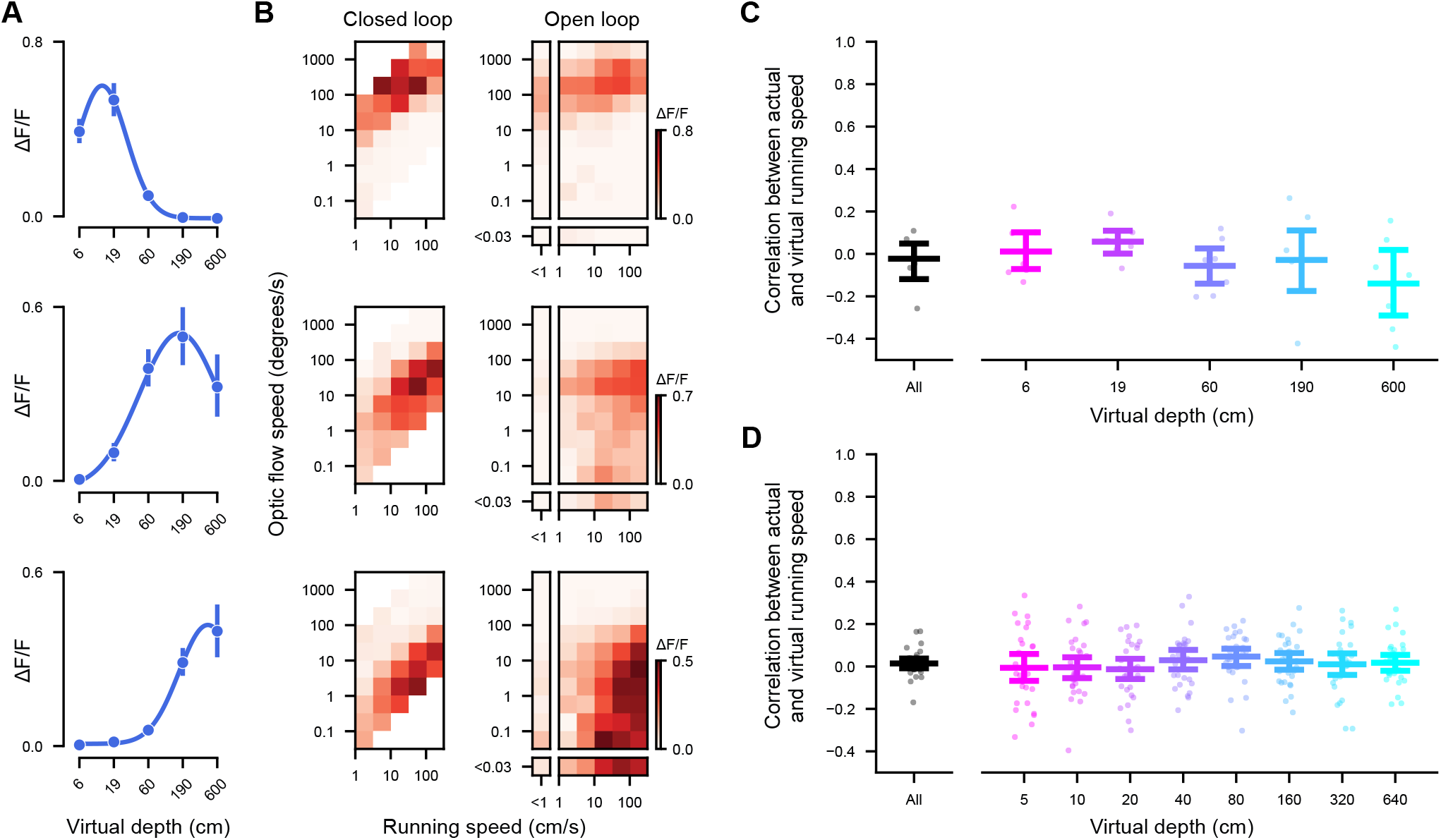
Optic flow and running speed tuning on open loop trials. **(A)** Virtual depth tuning of 3 example neurons. **(B)** Responses of the neurons in **A** as a function of both running and optic flow speeds in closed loop (left) and open loop (right) recordings. **(C-D)** Correlation coefficient between the actual and virtual running speeds during open loop recordings with 5 **(C)** or 8 **(D)** virtual depths for individual imaging sessions.

**Figure S6.**
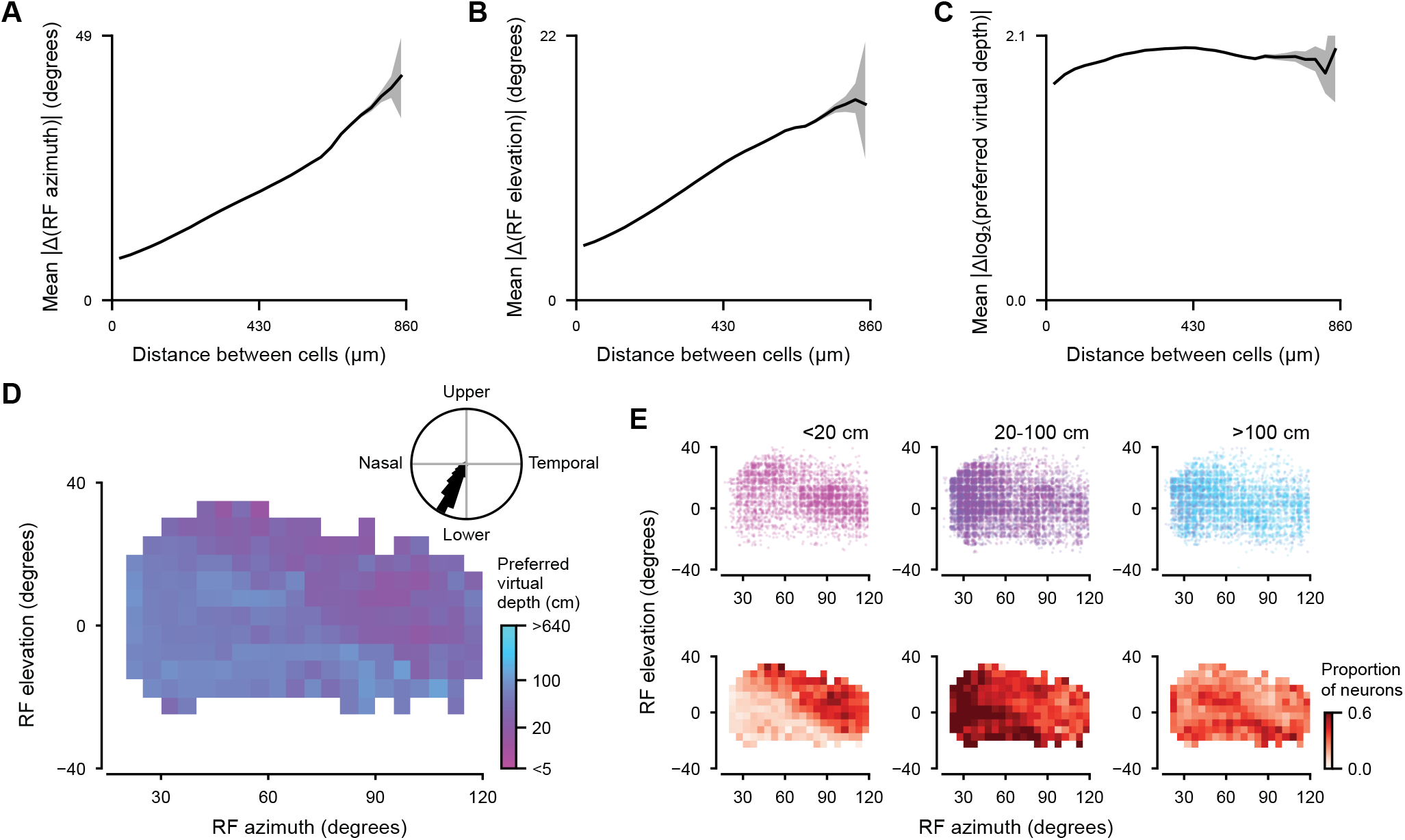
Distribution of depth preferences across the visual field. **(A-C)** Mean pairwise distance of RF azimuth **(A)**, RF elevation **(B)** or log-preferred depth **(C)** between depth-tuned neurons with significant RFs binned by the distance (N = 20,554 neurons from 85 sessions). Pairs of neurons within 10*µ*m from each other were excluded to avoid the effects of fluorescence contamination. Shading – 95% confidence interval computer by bootstrap resampling neuron pairs. **(D)** Median uncorrected preferred virtual depth as a function of RF location (N = 20,554 neurons from 85 sessions). Inset – direction of the gradient of preferred virtual depth across the visual field estimated across hierarchical bootstrap samples of the dataset. **(E)** Top – RF locations of depth-selective neurons tuned to near (<20 cm, N = 4,518 neurons), intermediate (20-100 cm, N = 10,325 neurons) and far (>100 cm, N = 5,711 neurons) virtual depths without correcting for RF location. Bottom – proportion of depth-selective neurons in each category as a function of RF location.

